# Invasion is accompanied by dietary contraction in Ponto-Caspian amphipods

**DOI:** 10.1101/2023.08.08.552405

**Authors:** Denis Copilaș-Ciocianu, Andrius Garbaras, Eglė Šidagytė-Copilas

**Affiliations:** Laboratory of Evolutionary Ecology of Hydrobionts, Nature Research Centre, Vilnius, Lithuania; Isotopic Research Laboratory, Center for Physical Sciences and Technology, Vilnius, Lithuania

**Keywords:** Crustacea, diet, functional morphology, invasive, native, niche, stable isotopes, range

## Abstract

A species’ expansion beyond the native range is often assumed to be associated with an increased dietary niche breadth. However, empirical evidence remains limited due to a scarcity of studies comparing both the parental and invaded ranges. Here, we test the trophic niche expansion hypothesis by examining stable isotopes and functional morphology across native (NW Black Sea) and invaded (SE Baltic Sea) ranges of two amphipods, *Dikerogammarus villosus* and *Pontogammarus robustoides*, originating from the Ponto-Caspian region – a major source of species invading Holarctic inland waters. Stable isotopes revealed that both species underwent a twofold contraction of the dietary niche with a shift towards decreased carnivory in the invaded range. This dietary shift was morphologically mirrored by an overall reduction of prey grasping appendages, antennae, and mouthpart palps. The magnitude of dietary and morphological change was greater in *D. villosus*. Our findings indicate that previous experimental reports of aggressive predation in *D. villosus* reflect opportunistic foraging and align with local stable isotope studies which generally indicate a low trophic position. We conclude that Ponto-Caspian species can undergo rapid, if non-intuitive, changes in both diet and functional morphology outside the native range, likely contributing to their invasive potential.

## 1. INTRODUCTION

Globalization-induced biotic homogenization has transformed ecosystems worldwide (Capinha et al., 2020; Herrick & Sarukhán, 2007; La Sorte et al., 2007; Shiganova, 2010) with alien and invasive species being one of the main threats to native biota (Blackburn et al., 2014; Pyšek et al., 2020). On the other hand, the arrival of alien species in new environments also poses unique opportunities to study rapid changes in ecological niches, adaptation, and eco-evolutionary dynamics (Miller et al., 2020; Moran & Alexander, 2014; Wiens et al., 2019). Understanding the extent to which invasive species can undergo rapid niche changes outside the native range is important for predicting their long-term persistence, adaptive potential, and future spread (Liu et al., 2020; Sotka et al., 2018).

Diet is one of the principal axes that defines an organism’s niche, which can be envisioned in the Hutchinsonian sense as an n-dimensional hypervolume (Carvalho & Cardoso, 2020; Hutchinson, 1957). A generalist diet is considered as one of the cornerstones underlying the success of invasive species as it enables capitalization on various food sources, depending on their availability (Shik & Dussutour, 2020; Snyder & Evans, 2006). Indeed, studies indicate that alien species tend to have broader trophic niches than their local resident counterparts (Feiner et al., 2013; Olsson et al., 2009; Rolla et al., 2020; Šidagytė et al., 2017; Sidorovich et al., 2010). However, at the intraspecific level comparatively much less is known about the extent to which the trophic niche of invasive populations has changed relative to native ones. Current evidence is rather limited, with studies indicating a wide pattern of variation, ranging from expansion to contraction or no shift at all (Balzani et al., 2021; Bissattini & Vignoli, 2017; Courant et al., 2017; Lancaster, 2020; Larson et al., 2010). This makes generalization difficult and emphasizes that a wide range of taxa need to be further studied (Balzani et al., 2021).

Globally, it is forecasted that arthropods will see the highest increase in the numbers of alien species by 2050, especially in North America and Europe (Seebens et al., 2021). This will incur additional ecological and economic costs in regions already facing one of the highest densities of invasive species worldwide (Diagne et al., 2021; Turbelin et al., 2017). A geographical area of particular interest in this context is the Ponto-Caspian region, which encompasses the Black, Azov, Caspian, and Aral Seas as well as the lower courses of their tributaries. This area is inhabited by a diverse endemic fauna of crustaceans, fishes, and mollusks with a peculiarly high tolerance to salinity fluctuations (Copilaș-Ciocianu & Sidorov, 2022; Paiva et al., 2018; Reid & Orlova, 2002). This ecological tolerance coupled with anthropogenic factors such as shipping, canal construction and intentional introductions has turned the Ponto-Caspian region into a major source of aquatic invaders throughout the Holarctic (Bij de Vaate et al., 2002; Copilaș-Ciocianu, Sidorov, et al., 2023; Soto et al., 2022). Invasion by Ponto-Caspian fauna is followed by a restructuring of communities and extinction of native species (Arbačiauskas & Gumuliauskaitė, 2007; Grabowski et al., 2009; Soto et al., 2022; van Kessel et al., 2016; Vanderploeg et al., 2002), sometimes with significant economic costs (Strayer, 2009). However, despite the broad geographical ranges and diversity of Ponto-Caspian taxa, comparative analyses of niche change between the native and invaded ranges are scarce and limited to climatic niches (Gallardo et al., 2013; Šidagytė-Copilas & Copilaș-Ciocianu, 2023). Changes in morphology have been reported in the round goby, but their ecological significance was not directly tested (Dashinov et al., 2020). Thus, it is not known whether the dietary niche of invasive populations of Ponto-Caspian taxa has changed in comparison to populations from the parental range.

Amphipod crustaceans are perhaps the most diverse group of Ponto-Caspian invaders, with up to 39 species representing 40% of the native fauna having spread beyond the native range (Copilaș-Ciocianu, Sidorov, et al., 2023). Of these, around a dozen species have substantially broad geographic distributions across the European continent and several species such as *Dikerogammarus villosus*, *D. haemobaphes,* and *Pontogammarus robustoides* are notoriously aggressive invaders with broad diets (Bacela-Spychalska & van der Velde, 2013; Berezina, 2007; Dick & Platvoet, 2000; Koester et al., 2016). Moreover, these invaders have negative effects on local native species (Arbačiauskas & Gumuliauskaitė, 2007; Dermott et al., 1998; Grabowski et al., 2009) via direct predation (Dick & Platvoet, 2000; Kinzler et al., 2009) or outcompeting them due to higher reproductive potential (Bacela & Konopacka, 2005; Grabowski, Bacela, et al., 2007; Pöckl, 2009). *Dikerogammarus villosus* has been nicknamed “the killer shrimp” and is considered one of the worst 100 invasive species worldwide (Rewicz et al., 2014). However, it remains unknown to what extent the trophic niche of these species has changed in the invaded range relative to the native one. Given the high ecomorphological and taxonomic diversity of gammaroid amphipods in the native Ponto-Caspian region (Copilaș-Ciocianu & Sidorov, 2022) it is likely that native dietary niches are narrower due to competition between numerous closely related sympatric species (Schluter, 2000). As such, we hypothesize that the invasive populations of Ponto-Caspian amphipods exhibit a broader trophic niche than native populations due to ecological release (Herrmann et al., 2021).

We tested this hypothesis on the widespread species *D. villosus* and *P. robustoides* by comparing invasive populations along the SE shore of the Baltic Sea (northern invaded range) to native source populations along the NW shore of the Black Sea. The invasion history of these species is well known and studied phylogeographically, confirming the NW Black Sea origin of alien Baltic populations (Copilaş-Ciocianu et al., 2022, 2023; Cristescu & Hebert, 2005; Jażdżewski, 1980; Rewicz et al., 2015). We employed a two-pronged approach that combines stable carbon and nitrogen isotopes and functional morphological traits related to prey detection (antennae and eyes), acquisition (gnathopods, i.e., grasping legs) and processing (mouthparts and stomach).

## 2. MATERIAL AND METHODS

### Study system

The two focal species, *D. villosus* and *P. robustoides*, are part of the Ponto-Caspian radiation of gammaroid amphipods, the second largest lacustrine amphipod radiation in the world, after Baikal Lake (Copilaș-Ciocianu, Palatov, et al., 2023; Copilaș-Ciocianu & Sidorov, 2022). Ponto-Caspian amphipods are an ecologically and morphologically diverse group that comprises four ecomorphs: crawlers associated with coarse substrate, diggers associated with fine substrate, clingers associated with plants, and symbionts. Our focal species belong to two ecomorphs with *D. villosus* being a crawler while *P. robustoides* is a digger (Copilaș-Ciocianu & Sidorov, 2022). As such they are both characterized by substantial morphological divergence and are thus good candidates for comparative studies.

These two species are also two of the most notorious Ponto-Caspian invaders, with extensively documented negative effects on recipient communities and extinction of native taxa (Arbačiauskas et al., 2017; Rewicz et al., 2014). Their distribution patterns and introduction modes differ substantially, with *D. villosus* being narrowly distributed in the native range (mainly NW and N Black Sea coast), but widespread in the invaded range (from Britain to W Russia and Latvia to Italy), where it invaded by natural dispersal and ballast water (Copilaș-Ciocianu, Sidorov, et al., 2023; Minchin et al., 2019; Rewicz et al., 2014). Conversely, *P. robustoides* is very broadly distributed in the native range (both Black and Caspian Seas) but has a narrower alien distribution. It spread by natural dispersal only in the middle to lower course of large rivers that drain into the Ponto-Caspian basin, and was intentionally introduced in the Baltic Sea region, initially in Lithuania (Arbačiauskas et al., 2017), from where it has spread throughout the entire SE coast (Copilaș-Ciocianu, Sidorov, et al., 2023; Copilaş-Ciocianu & Šidagytė-Copilas, 2022; Moedt & Van Haaren, 2018). Both species colonized the SE Baltic coast during the 1990s (Bij de Vaate et al., 2002; Grabowski, Jazdzewski, et al., 2007).

### Sampling

We sampled both species across their invaded range along the Baltic coast in 2020 at six sites (two each in Poland and Latvia, one each in Lithuania and Estonia) and native populations along the NW Black Sea coast in 2021 at seven sites (four in Ukraine, two in Romania and one in Bulgaria) (Fig. 1 and Table S1). All the localities were brackish transitional habitats to control for the potential effect of salinity on diet (Arbačiauskas et al., 2013). In both years animals were collected during late summer/early autumn (August-September) by kick-sampling with a hand net from all available habitats up to 1.5 m depth.

**Fig. 1.**
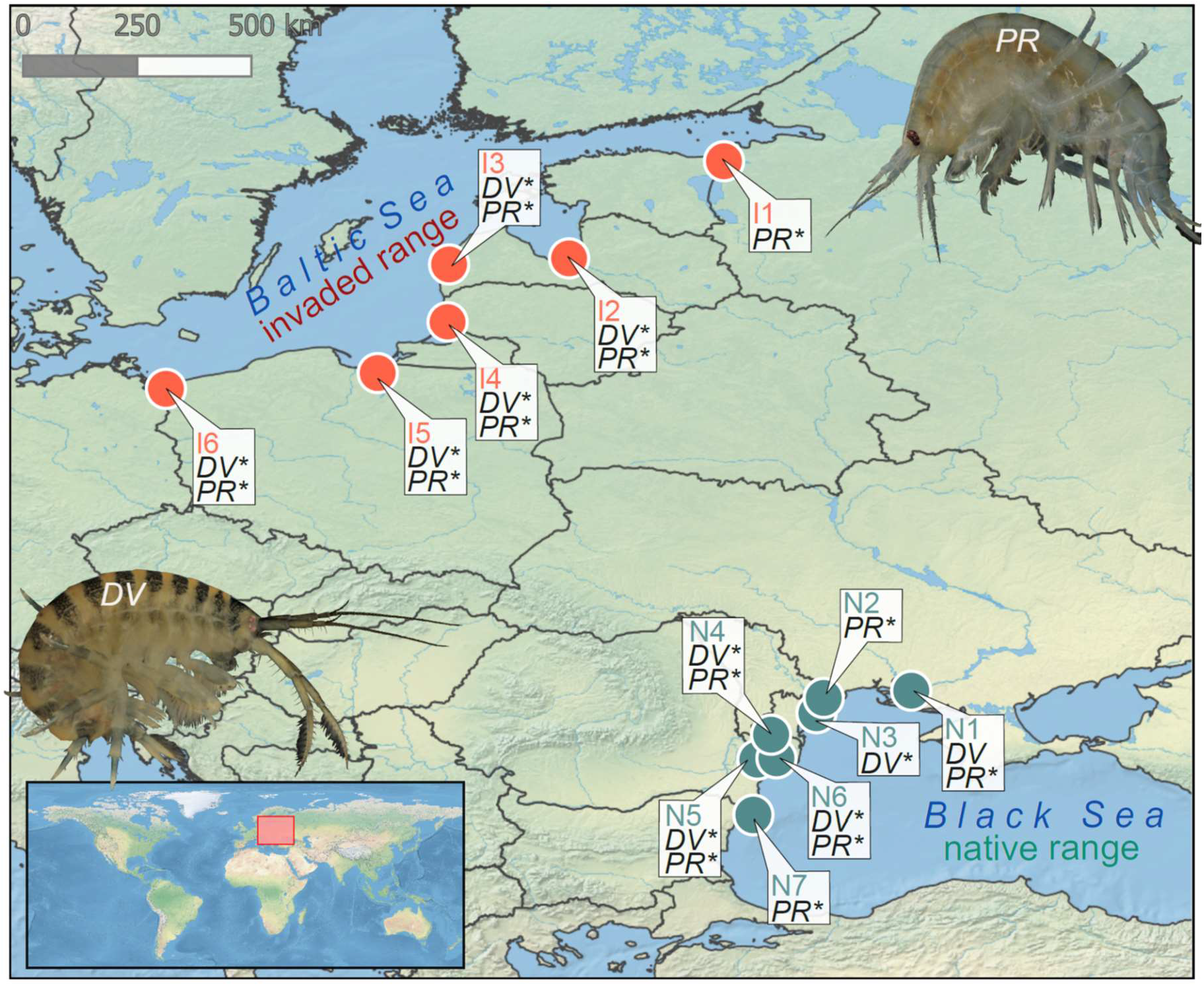
Sampling localities across the native (green points and site codes) and invaded (red points and site codes) ranges. Species abbreviations: DV – *Dikerogammarus villosus*, PR – *Pontogammarus robustoides*. Asterisks indicate populations analyzed for morphology and stable isotopes, while absence of asterisks indicates populations analyzed only for stable isotopes.

Largest available adult amphipods used for morphological study were preserved in 96% ethanol directly in the field. Amphipods of all available sizes for stable isotope analysis as well as mollusks or chironomids (intended as baseline organisms), were collected alive, transported in coolers, and kept overnight in filtered (0.7 µm) and aerated water from the site to empty the gut. The next day they were blotted and stored at –20°C until further treatment in the laboratory.

### Stable isotope analysis (SIA)

For SIA of C and N, the animals were defrosted at +4°C. Amphipods were identified using the latest keys (Copilaș-Ciocianu & Sidorov, 2022) and sexed, and their body size from the tip of the rostrum to the end of telson was measured under a stereomicroscope to the nearest 1 mm. Whole individual amphipods and the soft tissues of individual baseline mollusks were then dried at +60°C for 48 h, grinded to fine powder in an agate mortar, and weighted into tin capsules. In case of gravid females, the eggs were removed from the marsupium prior to drying. Each SIA sample represented a single individual unless (as in the case of the smallest juveniles) a single specimen was not enough for the target sample mass of 0.8–1.1 mg. Depending on specimen availability, we aimed to represent each available 1 mm group of a population by also including enough adult (> 7 mm) females and males as well as juveniles (< 7 mm). In total, 131 samples of *D. villosus* and 125 samples of *P. robustoides* were analyzed with a median of 21 samples per species/sex-juveniles/range group (range 11–34) and a median of 4 samples per sex-juveniles/site (range 0–9; juveniles were sometimes absent). Additionally, we considered a total of 52 samples of the gammarid *Obesogammarus crassus* to estimate the baseline isotope ratios (see below). See Table S1 for more details on sample numbers. The δ^13^C and δ^15^N measurements were conducted at the Isotopic Research Laboratory of the Centre for Physical Sciences and Technology in Vilnius, Lithuania. A Thermo Delta V isotope ratio mass spectrometer connected to Flash EA1112 elemental analyzer were used for isotope ratio mass spectrometry measurements with an overall precision of 0.1 ‰ for δ^13^C and 0.15 ‰ for δ^105^N. All SIA data are available in Figshare (doi will be provided during revision).

### Morphometric measurements

For morphometric analyses we conducted 39 measurements on functional morphological traits that directly reflect the diet of amphipods. The measurements are grouped as follows: 1) body length and head length as two complementary methods of body size (also used for standardization of subsequent measurements); 2) traits associated with food/prey detection – eye surface and lengths of antennary flagella and peduncles of both antenna pairs (8 measurements) (Caine, 1974; Copilaș-Ciocianu et al., 2021); 3) traits associated with food/prey grasping and handling – size and shape of both gnathopod pairs, i.e. grasping appendages (16 measurements) (Copilaș-Ciocianu et al., 2021; Premate et al., 2021); 4) traits associated with food/prey processing and digestion – lengths, surfaces, and meristic counts of mouthparts and stomach (13 measurements) (Coleman, 1991; Copilaș-Ciocianu et al., 2021;Hutchins et al., 2014; Mayer et al., 2009; Steele & Steele, 1993; Watling, 1993). A full list of measurements is provided in Table S2. We aimed at measuring ten adult specimens per sex per sampling site for each of the two species. In total we measured 356 individuals (166 *D. villosus* and 190 *P. robustoides*) with a median of 44 individuals (range 39–51) per species/sex/range and a median of 10 individuals (range 3–11) per sex/site (Table S1).

Prior to dissections, ethanol preserved animals were incubated overnight in a 2% lactic acid solution (LegoChem Biosciences Inc., Daejeon, South Korea) at room temperature and subsequently transferred for at least three hours to a 2:1 solution of 70% ethanol and glycerol (Carl Roth Gmbh & Co. Kg, Karlsruhe, Germany) to soften and partially clear the cuticle. Afterwards, animals were dissected in glycerol under a Nikon SMZ1000 stereomicroscope with the help of microsurgical Vannas spring scissors and 0.2 mm needles. Only the right-sided mouthparts were dissected as they are asymmetric in amphipods. The antennae and gnathopods were dissected from either side of the body, depending on the completeness of the specimen. All dissected appendages were glycerol-mounted on microscope slides and photographed under a Nikon SMZ1000 stereomicroscope or a Nikon Eclipse Si microscope equipped with a PixeLINK M15C-CYL camera and manufacturer provided software (PixeLINK Capture 3). Measurements were taken with Digimizer 4 (https://www.digimizer.com/). All raw measurements are available in Table S3 and Figshare (doi will be provided during revision).

### Stable isotope metrics of trophic position (TP)

Since the obtained data indicated that our samples had a varying C:N mass ratio often over the threshold of 3.5, prior to any analyses we corrected the measured δ^13^C values for lipid content using formulas for aquatic macroinvertebrates (Post et al., 2007). To standardize the trophic position of the focal amphipods for comparability between different sites, we calculated the trophic level (TL) and littoral reliance (LR) of the consumers using these formulas (Nilsson et al., 2012; Post, 2002; Zanden & Rasmussen, 2001):

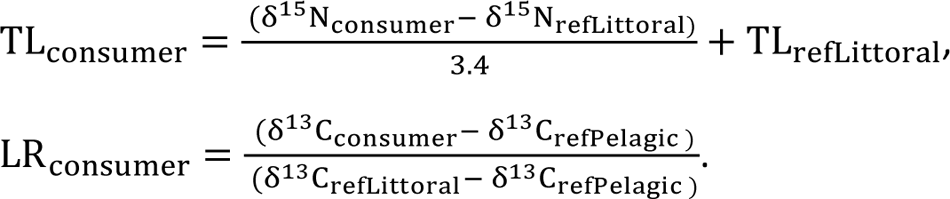

Calculating these indices required suitable baselines to reflect both the δ^13^C and δ^15^N of littoral primary consumers (refLittoral) and the δ^13^C of pelagic primary consumers (refPelagic). For the littoral references we intended to use grazing mollusks, such as *Radix* sp., however, in our dataset they were often way more ^13^C-depleted and sometimes more ^15^N-enriched than most of the focal amphipods, thus were not fit for the calculations. Instead, our data showed that another smaller species of Ponto-Caspian gammarid, namely *O. crassus*, available from both native and invaded ranges, was always the most ^13^C-enriched and ^15^N-depleted. Therefore, we resorted to considering its 2–3 mm juveniles as true herbivores (second trophic level) and littoral reference organisms. Since *O. crassus*, or only its juveniles, were not always available, we modelled their position in isotopic space of each site across the data of all three gammarids using mixed linear models of δ^13^C and δ^15^N as a function of Size (i.e., body length) and Species allowing for their interaction and a random intercept and slope for Size by Site (Size was centered by dataset mean). Species effect was statistically significant in both models (*P* ≤ 0.012), while the effect of Size was also marginally significant in the δ^15^N model (*P* = 0.07) (see conditional plots of main model effects in Fig. S1). Meanwhile, available filtering organisms (mostly mussel *Dreissena* sp.) were always the most ^13^C-depleted and thus their mean δ^13^C values were used as pelagic references. Information on all used baseline organisms and reference isotopic signatures is available in Table S1.

### Statistical testing

Within the TP data of each species, we tested the effects of Size (continuous, centered by mean value of the corresponding dataset), Sex (female/male) and Range (native/invaded) on TL and LR using linear mixed effects models (LMEMs) allowing for a random intercept for Site. As juveniles could not be assigned to sex, we doubled these analyses by dataset. At first, we tested the effects of all three factors (Size, Sex, and Range), including their second order interactions, only within the adult specimen sub datasets (adult LMEMs). Then, we tested the effects of Size and Range, and the interaction, within full species (adult and juveniles) datasets (all data LMEMs).

For the morphometry data, we first size-corrected the 38 traits by regressing them against body length (both log-transformed) and extracting the residuals. We then summarized the variation in each species’ morphology by running principal component analyses (PCAs) on the residuals of the 38 traits (based on correlation matrices). We used multivariate analyses of variance (MANOVAs) to test for effects of Sex, Range, and the interaction on general species morphology (all 38 size-corrected traits) and on each of three trait complexes separately: 1) prey detection – antennae and eyes, 2) prey grasping and handling – gnathopods, and 3) prey processing – mouthparts and stomach. By fitting LMEMs with random Site effect, we tested the effects of Sex, Range, and the interaction on the log-transformed body length and each individual size-corrected trait.

All analyses were conducted in the R environment. The LMEMs were fitted using the R packages *lme4* v. 1.1-32 and *lmerTest* v. 3.1-3 (Kuznetsova et al., 2017). The package *visreg* v. 2.7.0 was used to visualize the factor effects tested within the LMEMs (Breheny & Burchett, 2017). The PCAs were ran with the assistance from the packages *FactoMineR* v. 2.8 and *factoextra* 1.0.7 (Kassambra & Mundt, 2022; Lê et al., 2008).

### Inferring niche changes

To estimate the magnitude and patterns of niche differentiation between native and invaded ranges within each species we used the n-dimensional hypervolume approach applied to both stable isotopes-derived TP and morphology (Blonder et al., 2014). Since LMEMs revealed ontogenetic shifts in TL and/or LR in both species (see Results), we used TL and LR residuals from regressions against body length within each species’ dataset to construct the TP hypervolumes. TP hypervolumes were constructed using either all individuals within native or invaded range (adults and juveniles) or separately for each sex (excluding juveniles). Morphology hypervolumes were based on the first three PCA axes (see above), which cumulatively explained over 60% of the morphology variation in case of both species (see Table S4) and were constructed for each sex separately given the strong sexual dimorphism present in gammarids.

Hypervolume pairs, representing the native and invasive ranges, were constructed using the *hypervolume* v. 3.1.0. R package (Blonder et al., 2014, 2018, 2022). For each pair we calculated unique volumes, overlap, and the distance between centroids. Subsequently, the hypervolume pairs were used as input in the R package *BAT* v. 2.9.2 (Cardoso et al., 2015) to estimate the total niche change using the β_total_ diversity index (equal to 1 minus Jaccard similarity) which ranges from 0 (identical) to 1 (completely dissimilar). The total differentiation was then decomposed into the β_replacement_ index (indicating niche shift) and the β_richness_ index (indicating niche contraction/expansion) (Carvalho & Cardoso, 2020).

## 3. RESULTS

### Trophic position

Isotopic biplots indicated that invasive populations of both species tend to occupy lower trophic levels and rely more on littoral carbon input than the native populations, this tendency being more pronounced in *D. villosus* (Fig. 2).

**Fig. 2.**
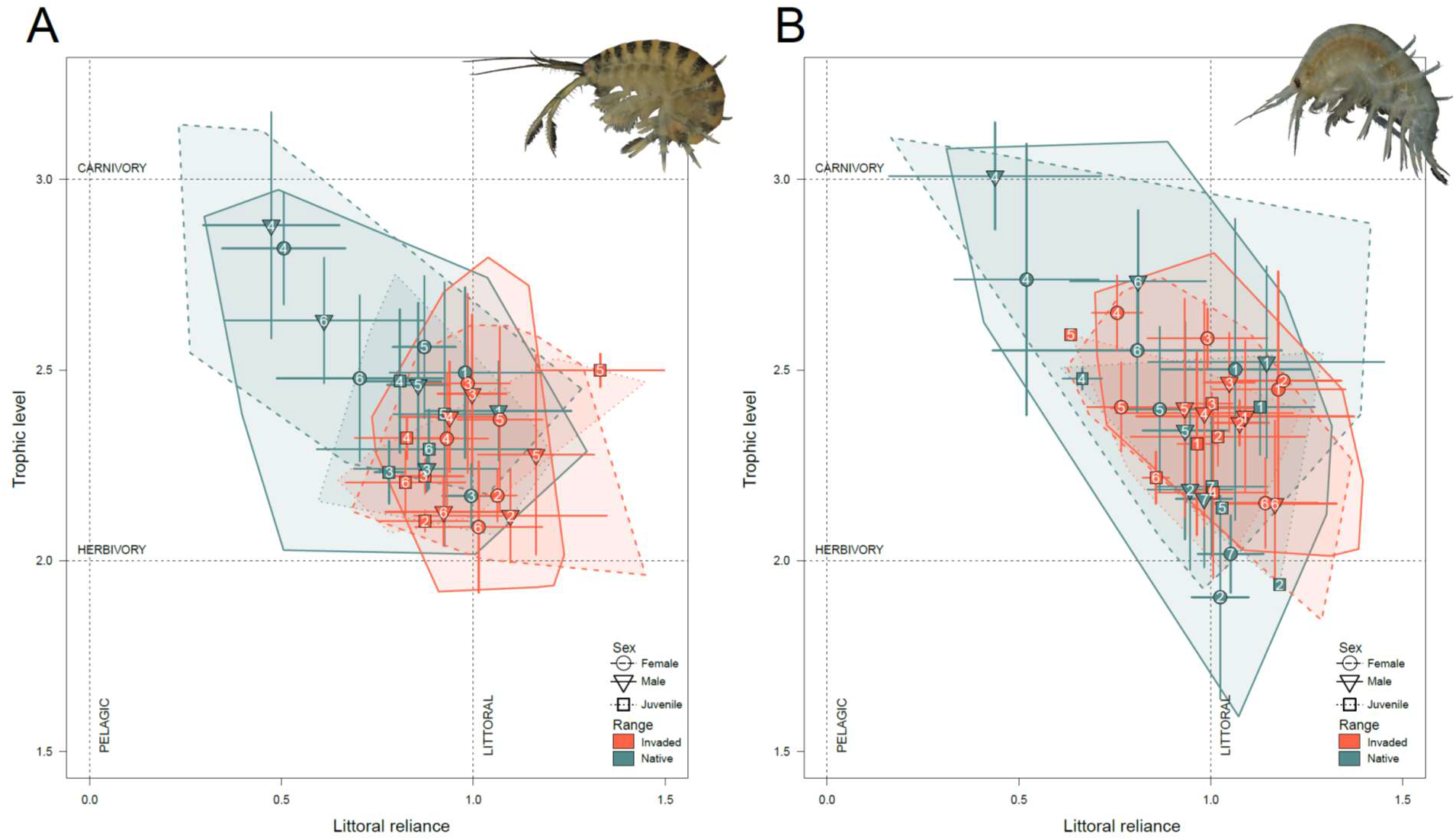
Standardized stable isotope biplots showing trophic positions of (A) *Dikerogammarus villosus* and (B) *Pontogammarus robustoides* with convex hulls indicating differences between native and the invaded ranges by sex group. Points and whiskers represent means and standard deviations per site/sex group. Numbers correspond to populations in Fig. 1.

Both LMEMs (either adults only or combined adults and juveniles) of *D. villosus* TL indicated significantly steeper ontogenetic slopes in the invaded range (Size:Range effects: *F* ≥ 4.5, *P* ≤ 0.04) (Table 1, Fig. S2). The combined adult and juvenile LMEM of LR for this species also indicated a change in ontogenetic slope from negative in the native to positive in the invaded range (Size:Range effect: *F* = 4.7, *P* = 0.03), reflected only by a marginally significant range effect in the adult LMEM (Range effect: *F* = 5.1, *P* = 0.05) (Table 1, Fig. S3). Neither LMEM for this species indicated a significant differentiation by sex.

**Table 1.**
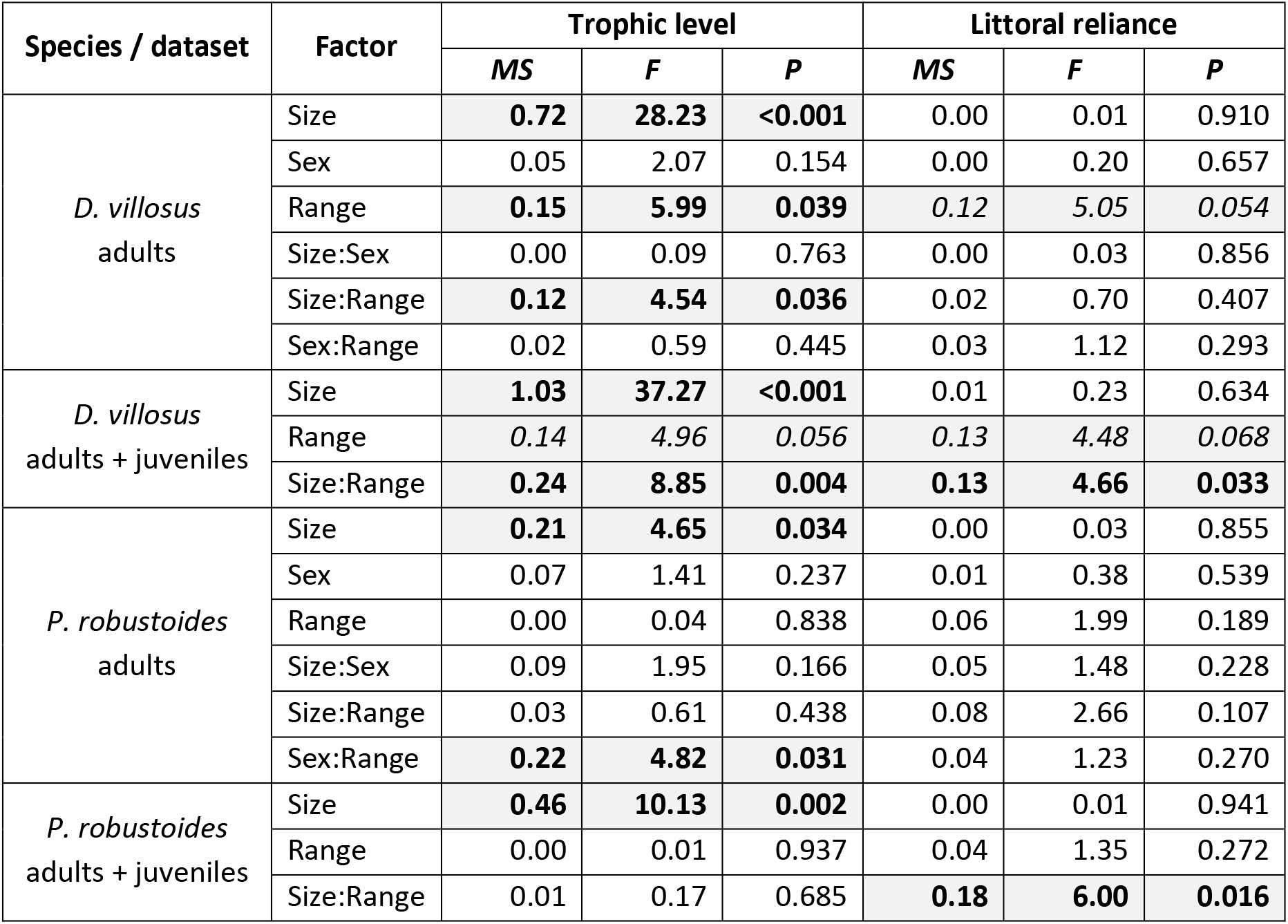
Results of linear mixed effects models (type III analysis of variance with Satterthwaite’s method for approximating degrees of freedom) testing the effects of Size, Sex (female/male) and Range (native/invaded) on stable isotope indices of trophic position (trophic level and littoral reliance) within adult or adults plus juveniles datasets of *Dikerogammarus villosus* and *Pontogammarus robustoides*. Significant effects (*P* < 0.05) are in bold and marginally significant results (*P* < 0.1) are in italic font; both are highlighted. See Figs. S2–S3 for visualization of model effects.

| Species / dataset | Factor | Trophic level |  |  | Littoral reliance |  |  |
| --- | --- | --- | --- | --- | --- | --- | --- |
|  |  | *MS* | *F* | *P* | *MS* | *F* | *P* |
| *D. villosus*<br>adults | Size | **0.72** | **28.23** | **<0.001** | 0.00 | 0.01 | 0.910 |
|  | Sex | 0.05 | 2.07 | 0.154 | 0.00 | 0.20 | 0.657 |
|  | Range | **0.15** | **5.99** | **0.039** | *0.12* | *5.05* | *0.054* |
|  | Size:Sex | 0.00 | 0.09 | 0.763 | 0.00 | 0.03 | 0.856 |
|  | Size:Range | **0.12** | **4.54** | **0.036** | 0.02 | 0.70 | 0.407 |
|  | Sex:Range | 0.02 | 0.59 | 0.445 | 0.03 | 1.12 | 0.293 |
| *D. villosus*<br>adults + juveniles | Size | **1.03** | **37.27** | **<0.001** | 0.01 | 0.23 | 0.634 |
|  | Range | *0.14* | *4.96* | *0.056* | *0.13* | *4.48* | *0.068* |
|  | Size:Range | **0.24** | **8.85** | **0.004** | **0.13** | **4.66** | **0.033** |
| *P. robustoides*<br>adults | Size | **0.21** | **4.65** | **0.034** | 0.00 | 0.03 | 0.855 |
|  | Sex | 0.07 | 1.41 | 0.237 | 0.01 | 0.38 | 0.539 |
|  | Range | 0.00 | 0.04 | 0.838 | 0.06 | 1.99 | 0.189 |
|  | Size:Sex | 0.09 | 1.95 | 0.166 | 0.05 | 1.48 | 0.228 |
|  | Size:Range | 0.03 | 0.61 | 0.438 | 0.08 | 2.66 | 0.107 |
|  | Sex:Range | **0.22** | **4.82** | **0.031** | 0.04 | 1.23 | 0.270 |
| *P. robustoides*<br>adults + juveniles | Size | **0.46** | **10.13** | **0.002** | 0.00 | 0.01 | 0.941 |
|  | Range | 0.00 | 0.01 | 0.937 | 0.04 | 1.35 | 0.272 |
|  | Size:Range | 0.01 | 0.17 | 0.685 | **0.18** | **6.00** | **0.016** |

For *P. robustoides*, only the adult data LMEM of TL indicated differentiation by range as females tended to occupy lower TL in the invaded than in the native range, while the TL of males appeared to remain similar (Sex:Range effect: *F* = 4.8, *P* = 0.03) (Table 1, Fig. S2). Similarly to *D. villosus*, for *P. robustoides* the combined adult and juvenile LMEM of LR revealed a significant change of ontogenetic slope from negative in the native to positive in the invaded range (Size:Range effect: *F* = 6.0, *P* = 0.02) (Table 1, Fig. S3), although this pattern was not statistically significant in the adult data LMEM (Size:Range effect: *F* = 2.7, *P* = 0.11).

### Morphology

The first two axes of the PCAs (Fig. 3) revealed tendencies that the morphology of both species has shifted in a similar fashion in the invaded range, with a general decrease in the size of gnathopod propodi, lengths of antennal segments as well as mandibular and maxillipedal palps. The overall magnitude of change was greater in *D. villosus*. MANOVA tests using either all traits or each of the three main trait complexes (prey detection, grasping, or processing) independently revealed a highly significant differentiation (Table 2). The interaction between sex and range was significant (*F* ≥ 2.0, *P* ≤ 0.04) in all cases except for the food detection trait complex in *D. villosus* and food processing trait complex in *P. robustoides* (*F* ≤ 1.0, *P* ≥ 0.5). Body length, as revealed by LMEMs, was significantly greater in the invaded range (*F* = 29.7, *P* = 0.001) in *D. villosus*, but not in *P. robustoides* (*F* = 0.5, *P* = 0.5) (Fig. 4). Results of the LMEMs for each individual trait are presented in Table S5 and Figs. S4–S5.

**Fig. 3.**
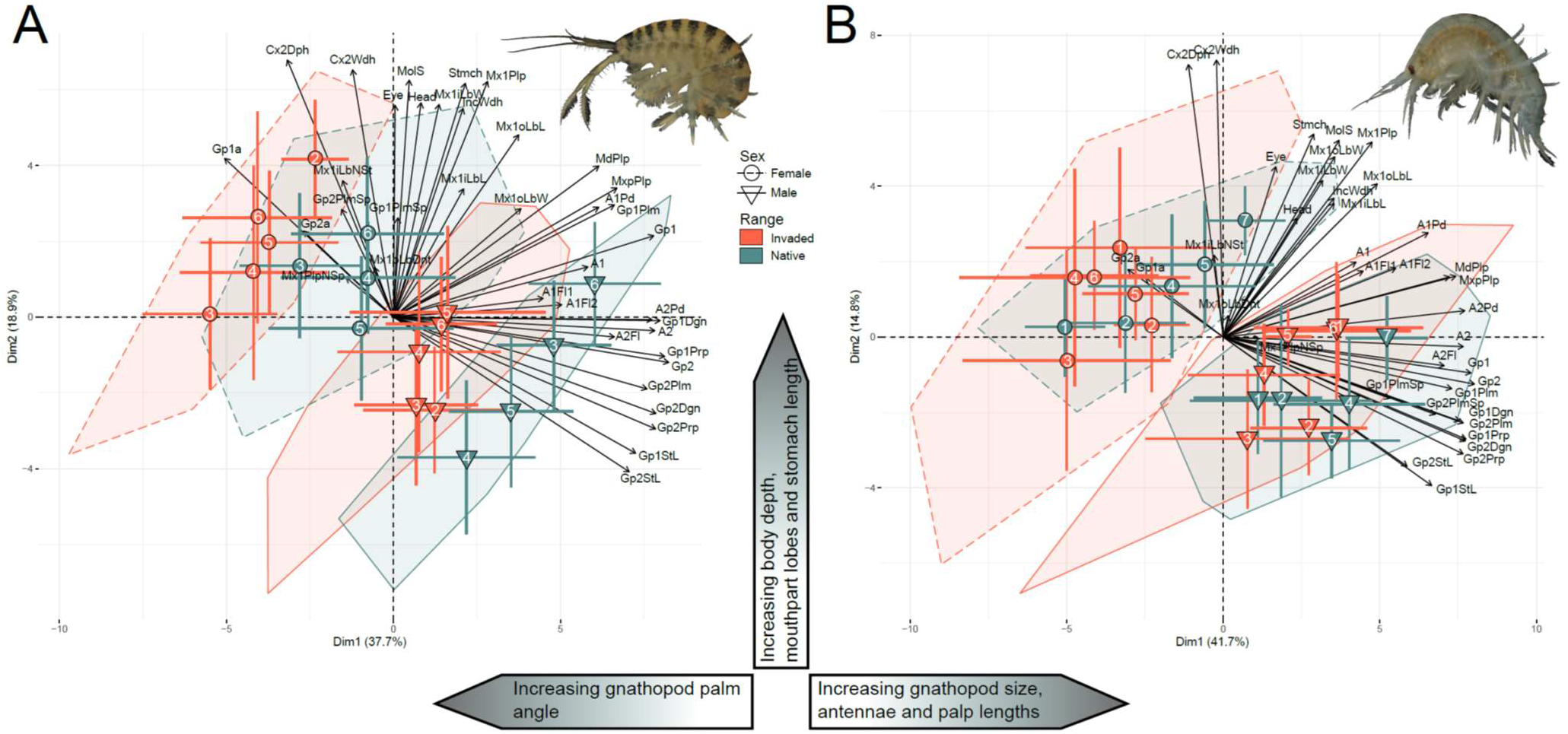
Biplots of the two first axes from the principal component analyses of morphological traits in (A) *Dikerogammarus villosus* and (B) *Pontogammarus robustoides* with convex hulls indicating differences between native and the invaded ranges by sex group. Points and whiskers represent means and standard deviations per site/sex group (see Table S2 for trait definitions and Table S4 for their contributions to the axes). Numbers correspond to populations in **Fig. 1**.

**Fig. 4.**
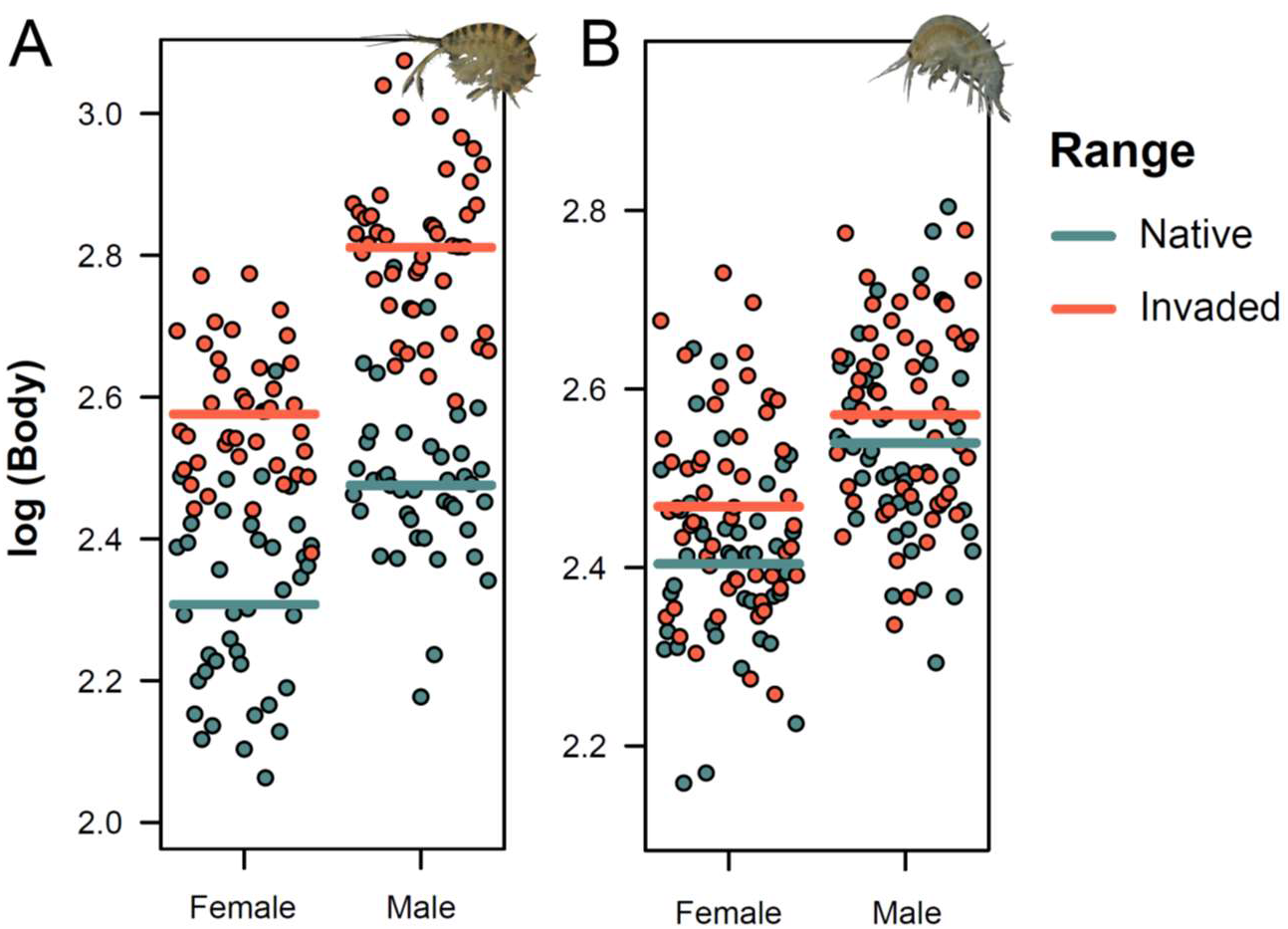
Sex and Range effects on body size in (A) *Dikerogammarus villosus* and (B) *Pontogammarus robustoides* – visualization of linear mixed effects models (see Table S5 for additional information).

**Table 2.** Results of MANOVAs testing the effects of Sex (female/male) and Range (native/invaded) on all analyzed morphometric traits or each of the three trait complexes in *Dikerogammarus villosus* and *Pontogammarus robustoides*. Significant effects (*P* < 0.05) are in bold and highlighted.

| Species | Traits | Factor | Pillai-Bartlett | F | P |
| --- | --- | --- | --- | --- | --- |
| *D. villosus* | All | Sex | 0.92 | 36.89 | **<0.001** |
|  |  | Range | 0.74 | 9.17 | **<0.001** |
|  |  | Sex:Range | 0.38 | 1.98 | **0.003** |
|  | Prey detection<br>(eyes + antennae) | Sex | 0.67 | 39.87 | **<0.001** |
|  |  | Range | 0.39 | 12.55 | **<0.001** |
|  |  | Sex:Range | 0.05 | 0.92 | 0.505 |
|  | Prey handling<br>(gnathopods) | Sex | 0.89 | 84.77 | **<0.001** |
|  |  | Range | 0.60 | 16.15 | **<0.001** |
|  |  | Sex:Range | 0.15 | 1.84 | **0.037** |
|  | Prey processing<br>(mouthparts + stomach) | Sex | 0.62 | 18.52 | **<0.001** |
|  |  | Range | 0.34 | 5.99 | **<0.001** |
|  |  | Sex:Range | 0.19 | 2.78 | **0.001** |
| *P. robustoides* | All | Sex | 0.86 | 23.70 | **<0.001** |
|  |  | Range | 0.72 | 9.87 | **<0.001** |
|  |  | Sex:Range | 0.35 | 2.07 | **0.001** |
|  | Prey detection<br>(eyes + antennae) | Sex | 0.60 | 33.96 | **<0.001** |
|  |  | Range | 0.39 | 14.04 | **<0.001** |
|  |  | Sex:Range | 0.14 | 3.72 | **<0.001** |
|  | Prey handling<br>(gnathopods) | Sex | 0.82 | 56.90 | **<0.001** |
|  |  | Range | 0.44 | 9.69 | **<0.001** |
|  |  | Sex:Range | 0.14 | 1.95 | **0.024** |
|  | Prey processing<br>(mouthparts + stomach) | Sex | 0.54 | 15.44 | **<0.001** |
|  |  | Range | 0.26 | 4.63 | **<0.001** |
|  |  | Sex:Range | 0.07 | 0.99 | 0.465 |

### Hypervolumes

With respect to trophic niche, the hypervolume analyses retrieved consistent patterns regardless of species, sex, or life stages combination. There was a moderately high degree of general differentiation among ranges in both *D. villosus* (β_total_ = 0.66–0.61) and *P. robustoides* (β_total_ = 0.55–0.65) (Table 3). In both species there are clear indications of a trophic niche contraction since the contribution of the β_richness_ index to the total differentiation (β_total_) ranged from 64 to 84% in *D. villosus* and from 89 to 93% in *P. robustoides* and the invaded niche volume was around two-fold smaller than the native volume in both species (Table 3, Figs. S6–S11). Although generally weak, the remainder trophic shift tendency in DV could be noted.

**Table 3.**
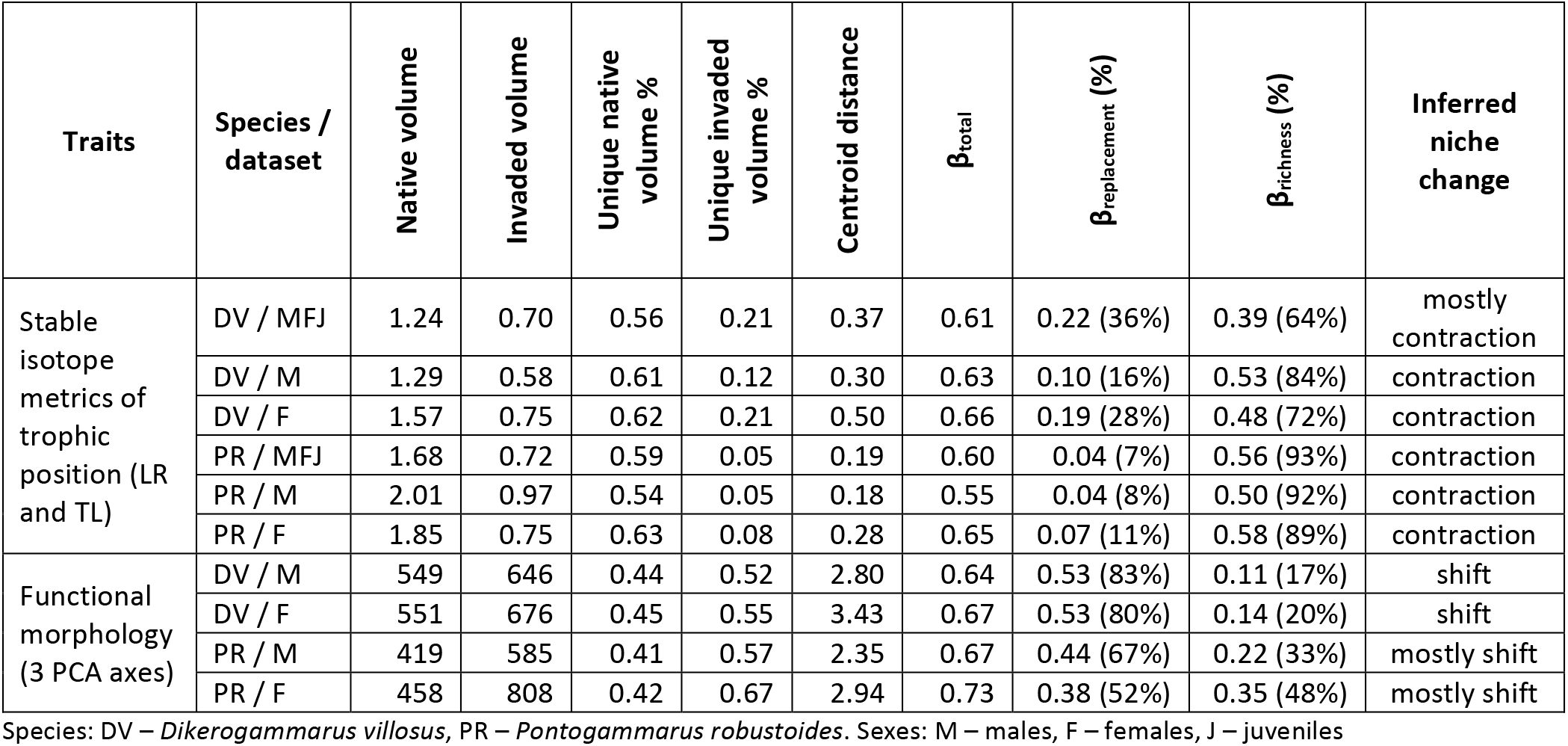
Results of hypervolume analysis of niche change patterns between native and invaded ranges in *Dikerogammarus villosus* (DV) and *Pontogammarus robustoides* (PR) by different traits and datasets: volume sizes, unique (unaffected by the overlap) part of each relatively to the volume size of each, distance between centroids, total dissimilarity and its partitioning into replacement (shift) and richness (contraction/expansion).

| Traits | Species / dataset | Native volume | Invaded volume | Unique native volume % | Unique invaded volume % | Centroid distance | $\beta_{\text{total}}$ | $\beta_{\text{replacement}}$ (%) | $\beta_{\text{richness}}$ (%) | Inferred niche change |
| --- | --- | --- | --- | --- | --- | --- | --- | --- | --- | --- |
| Stable isotope metrics of trophic position (LR and TL) | DV / MFJ | 1.24 | 0.70 | 0.56 | 0.21 | 0.37 | 0.61 | 0.22 (36%) | 0.39 (64%) | mostly contraction |
|  | DV / M | 1.29 | 0.58 | 0.61 | 0.12 | 0.30 | 0.63 | 0.10 (16%) | 0.53 (84%) | contraction |
|  | DV / F | 1.57 | 0.75 | 0.62 | 0.21 | 0.50 | 0.66 | 0.19 (28%) | 0.48 (72%) | contraction |
|  | PR / MFJ | 1.68 | 0.72 | 0.59 | 0.05 | 0.19 | 0.60 | 0.04 (7%) | 0.56 (93%) | contraction |
|  | PR / M | 2.01 | 0.97 | 0.54 | 0.05 | 0.18 | 0.55 | 0.04 (8%) | 0.50 (92%) | contraction |
|  | PR / F | 1.85 | 0.75 | 0.63 | 0.08 | 0.28 | 0.65 | 0.07 (11%) | 0.58 (89%) | contraction |
| Functional morphology (3 PCA axes) | DV / M | 549 | 646 | 0.44 | 0.52 | 2.80 | 0.64 | 0.53 (83%) | 0.11 (17%) | shift |
|  | DV / F | 551 | 676 | 0.45 | 0.55 | 3.43 | 0.67 | 0.53 (80%) | 0.14 (20%) | shift |
|  | PR / M | 419 | 585 | 0.41 | 0.57 | 2.35 | 0.67 | 0.44 (67%) | 0.22 (33%) | mostly shift |
|  | PR / F | 458 | 808 | 0.42 | 0.67 | 2.94 | 0.73 | 0.38 (52%) | 0.35 (48%) | mostly shift |

Species: DV – *Dikerogammarus villosus*, PR – *Pontogammarus robustoides*. Sexes: M – males, F – females, J – juveniles

Regarding morphology, the hypervolumes also revealed similar patterns irrespective of species and sex. The degree of general differentiation among ranges was substantial in both *D. villosus* (males: β_total_ = 0.64; females: β_total_ = 0.67) and *P. robustoides* (males: β_total_ = 0.67; females: β_total_ = 0.73) (Table 3).

Primarily, a shift in morphology can be inferred for both species and sexes because the β_replacement_ index had a higher contribution to the total differentiation than the β_richness_ index (80–83% in *D. villosus*; 52– 67% in *P. robustoides*). The volumes increased in the invaded range 1.2 times in *D. villosus* and 1.5–1.8 times in *P. robustoides*, also indicating signs of increased variation in morphology of the latter species (Table 3, Figs. S12–15).

## 4. DISCUSSION

Our study revealed that invasion of Ponto-Caspian amphipods is associated with a significant trophic niche contraction and a general shift towards decreased carnivory. Importantly, this shift was also detected morphologically as we observed an overall attenuation of traits associated with predation. These concordant changes, although counter to our dietary niche expansion hypothesis, highlight the rapid adaptive potential of Ponto-Caspian species.

Both *D. villosus* and *P. robustoides* have undergone a significant trophic niche contraction throughout the invaded SE Baltic range. Specifically, the volume of the invaded niche of both species is on average twice as small as the volume of the native niche, indicating a greater dietary plasticity in the parental range. Indeed, throughout the invaded Baltic range, both species have less predatory diets and rely more on littoral carbon sources, this being more obvious in *D. villosus*. This dietary shift is further reinforced by functional morphology where key traits that facilitate predation, such as relatively large gnathopod propodi as well as long antennae and mouthpart palps (Caine, 1974; Coleman, 1991; Copilaș-Ciocianu et al., 2021; Hutchins et al., 2014; Premate et al., 2021), are diminished in the invaded range.

Noteworthy, the magnitude of morphological change is greater in *D. villosus*, mirroring its more pronounced shift towards a lower trophic level. This is quite unexpected given that its body size has significantly increased in the invaded range, potentially opening a wider range of foraging options. Thus, the concordance between trophic niche and functional morphology across both species strongly supports a dietary niche shift towards decreased carnivory in the invaded range. This indicates that the often-observed morphological divergence among native and invasive populations of various taxa may indeed be ecologically relevant (Dashinov et al., 2020; Korablev et al., 2017; Le Gros et al., 2016; Zou et al., 2007).

Our findings of decreased predation outside the parental range seem contradictory to the experimental reports that *D. villosus* is a highly aggressive predator (Bacela-Spychalska & van der Velde, 2013; Dick & Platvoet, 2000; Kinzler et al., 2009), but are in accordance with local stable isotope-based field studies which generally suggest a low trophic position that is very similar to that of other co-occurring gammarids (Hellmann et al., 2015; Koester et al., 2016; Koester & Gergs, 2014; Sahm et al., 2021; Van Riel et al., 2006), but not always (Bacela-Spychalska & van der Velde, 2013; Mancini et al., 2021). However, given that stable isotopes reflect an individual’s long-long term diet (Araújo et al., 2007), this discrepancy might be explained by the opportunistic predatory behavior commonly encountered across Amphipoda (Dauby et al., 2001; Guerra-García & Tierno de Figueroa, 2009) and is supported experimentally in *D. villosus* when presented with multiple food choices (Médoc et al., 2018). Thus, it appears that aggressive predation may not be the main mechanism by which invasive gammarids replace native species in the wild.

Although we uncovered congruent dietary and morphological changes in both *D. villosus* and *P. robustoides*, the magnitude of these changes was greater in the former. An explanation for this is that *D. villosus* is ecologically more specialized than *P. robustoides*, and, thus, it might experience a stronger selective pressure in new environments. There are several lines of evidence for this. At the microhabitat scale *D. villosus* occurs predominantly on stony substrates in shallow water (Borza et al., 2018; Copilaș-Ciocianu & Sidorov, 2022; Kley & Maier, 2005; Poznańska-Kakareko et al., 2021), while *P. robustoides* has a very broad substrate preference occurring on fine substrates, coarse gravel, stones and even plants, and was reported at up to 200 m depth in the Caspian Sea (Copilaș-Ciocianu & Sidorov, 2022; Jermacz et al., 2015; Poznańska-Kakareko et al., 2021). At a large spatial scale *D. villosus* has a much narrower geographical distribution in the native range, being limited to the N-NW coast of the Black and Azov Seas, while *P. robustoides* is perhaps the most common species across the entire Ponto-Caspian basin (Copilaș-Ciocianu, Sidorov, et al., 2023). This clearly indicates that overall *P. robustoides* is ecologically more tolerant than *D. villosus* and therefore, might adapt or acclimate to novel conditions without undergoing pronounced dietary and morphological changes.

Our study adds to the growing body of evidence highlighting the diversity of patterns of dietary niche change that accompany biological invasions. Dietary niche contraction is not rare in invasive species when encountering resident competitors, thus it is often site or region specific (Balzani et al., 2021; Bissattini & Vignoli, 2017; Courant et al., 2017; Jackson et al., 2016). The contraction observed in Ponto-Caspian amphipods is contrary to our initial expectations of dietary expansion. We find this unexpected given that the investigated non-native waterbodies have a lower number of co-occurring amphipod species than in the native range (Arbačiauskas, 2008; Cărăușu et al., 1955; Copilaş-Ciocianu & Šidagytė-Copilas, 2022; Grabowski, Jazdzewski, et al., 2007; Martynov, 1924; Meßner & Zettler, 2018), where they evolved in situ since the Miocene (Copilaş-Ciocianu et al., 2020; Hou et al., 2014). Thus, one would expect a decreased competition and larger niche breadths in the invaded range and increased competition accompanied by a greater niche compaction in the native range due to many closely related co-evolved sympatric species (Rundell & Price, 2009; Simões et al., 2016). Following the ecological release hypothesis, this would translate either in a niche expansion or shift once the invasive species expand beyond the confines of the native range (Herrmann et al., 2021). However, we nevertheless observe a contraction of the dietary niche. The reasons for this are yet unclear but might be related to more intense pressure from predators (Scoleri et al., 2023), increased competition form more distantly related taxa (Wang et al., 2021) or conspecifics (Parent et al., 2014), or might even reflect a higher food abundance in the invaded range (Steinmetz et al., 2021).

Considering that both *D. villosus* and *P. robustoides* were first reported outside the native range in the first half of the 20^th^ century and both invaded the SE Baltic shores during the 1990s (Bij de Vaate et al., 2002; Grabowski, Jazdzewski, et al., 2007), it can be concluded that the observed dietary and morphological changes occurred in less than a century, possibly even a few decades. However, it remains to be confirmed whether these changes are true genetic adaptations or represent phenotypic plasticity (Santi et al., 2020; Vandepitte et al., 2014). Nevertheless, these rapid changes occurring concordantly in diet and morphology over decadal timespans highlight the adaptive potential of invasive Ponto-Caspian species (Davidson et al., 2011).

Future studies are needed to better understand the patterns uncovered herein. Common garden experiments could help identify whether the uncovered divergence among ranges results from adaptive evolution or plasticity. Furthermore, a greater number of populations needs to be studied to understand whether the uncovered differences represent the extreme ends of a gradient and to what extent freshwater habitats (not examined here) can further facilitate divergence.

## 5. CONCLUSIONS

We conclude that stable isotopes and functional morphology consistently indicate that the two widespread invasive amphipods *Dikerogammarus villosus* and *Pontogammarus robustoides* underwent a significant dietary contraction with a shift towards decreased predation in the invaded range. These changes occurred over decadal timespans and were more pronounced in *D. villosus*, suggesting an increased adaptive potential. Our study is the first to highlight rapid and concordant dietary and morphological changes in species originating from the Ponto-Caspian region – one of the most important global sources of inland aquatic invaders.

## AUTHOR CONTRIBUTIONS

Conceptualization: DCC and ESC; Data acquisition: DCC, ESC, AG; Data analysis: ESC, DCC; Writing: DCC, ESC.

## ACKNOWLEDGEMENTS

This study was financed by the Research Council of Lithuania (Contract No. S-MIP-20-26). We are grateful to Halyna Morhun and Mikhail Son for their invaluable help during the fieldwork in Ukraine.

## DATA AVAILABILITY STATEMENT

The data that support the findings of this study are available in the supplementary information and in Figshare (doi will be provided during the revision).

