## Supplementary material for "Invasion is accompanied by dietary contraction in Ponto-Caspian amphipods": Figure S

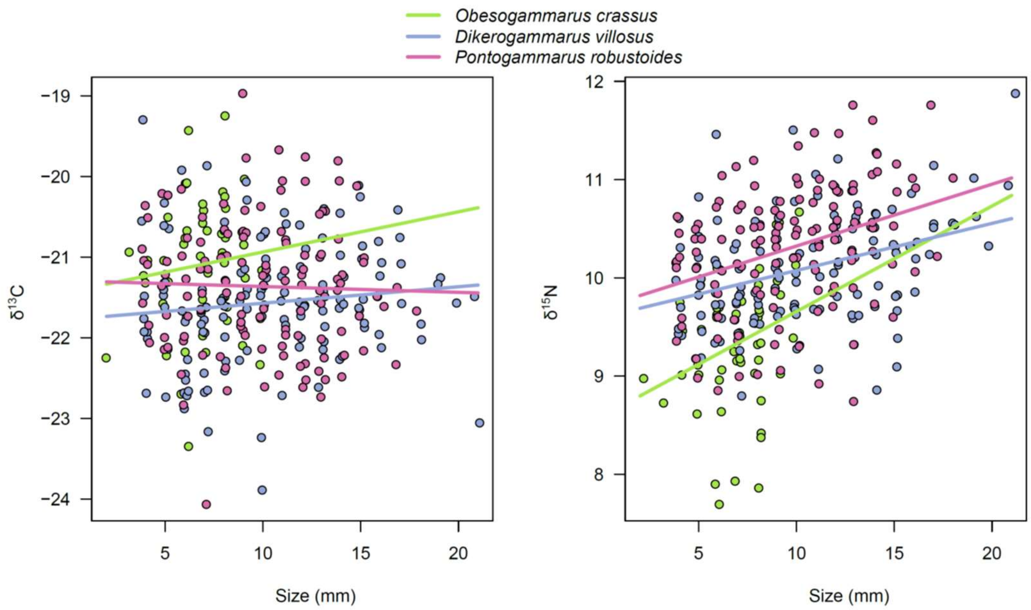


Fig. S1. Species and Size effects on $\delta^{15}N$ and $\delta^{13}C$ in three Ponto-Caspian amphipods – visualisation of linear mixed effects models used to predict the littoral reference values (see *2. Materials and Methods: Data analysis: Stable isotope metrics of trophic position* for notes on the models). Note that 2–3 mm juveniles of *O. crassus* tend to be the most $^{15}N$-depleted and $^{13}C$-enriched among studied species.

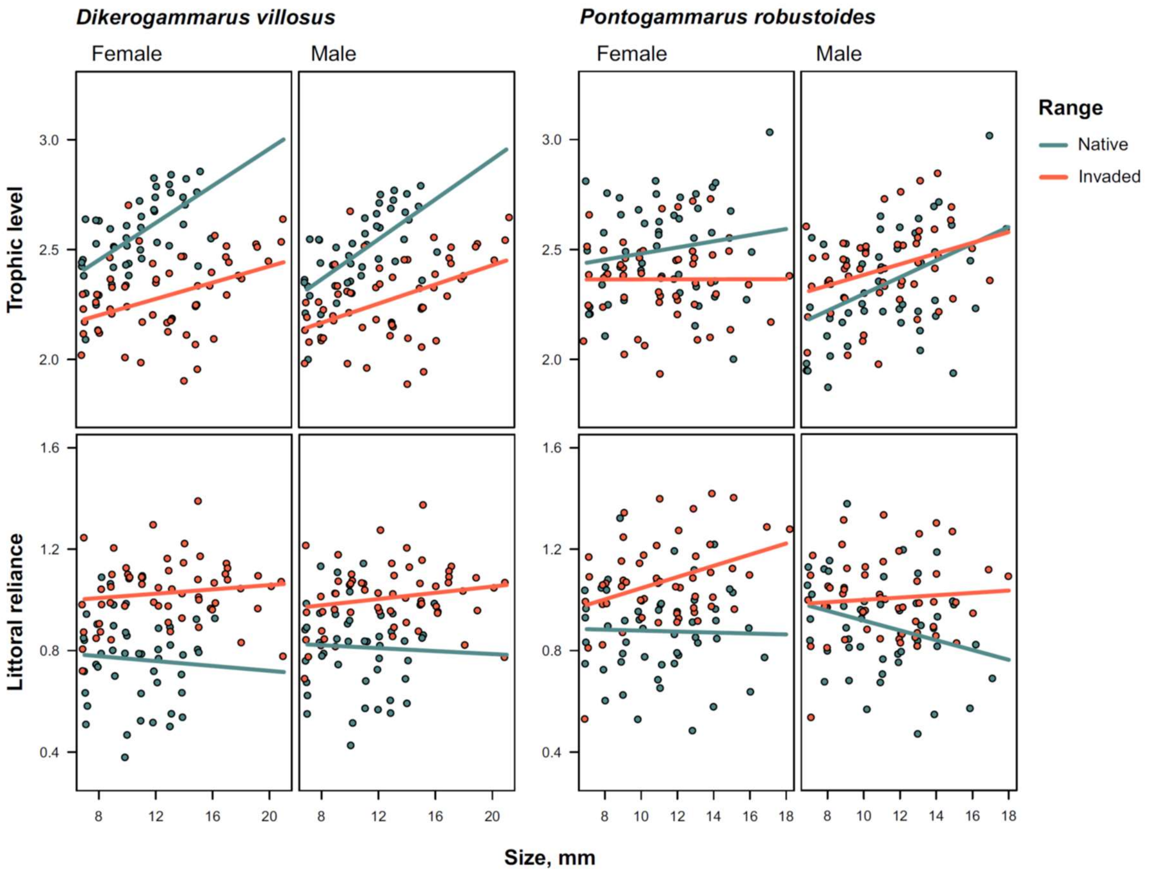


Fig. S2. Size, Sex, and Range effects on stable isotope metrics of trophic position – trophic level and littoral reliance – in two Ponto-Caspian amphipods – visualisation of linear mixed effects models within adult specimen subdatasets (see Table 1 for effect tests).

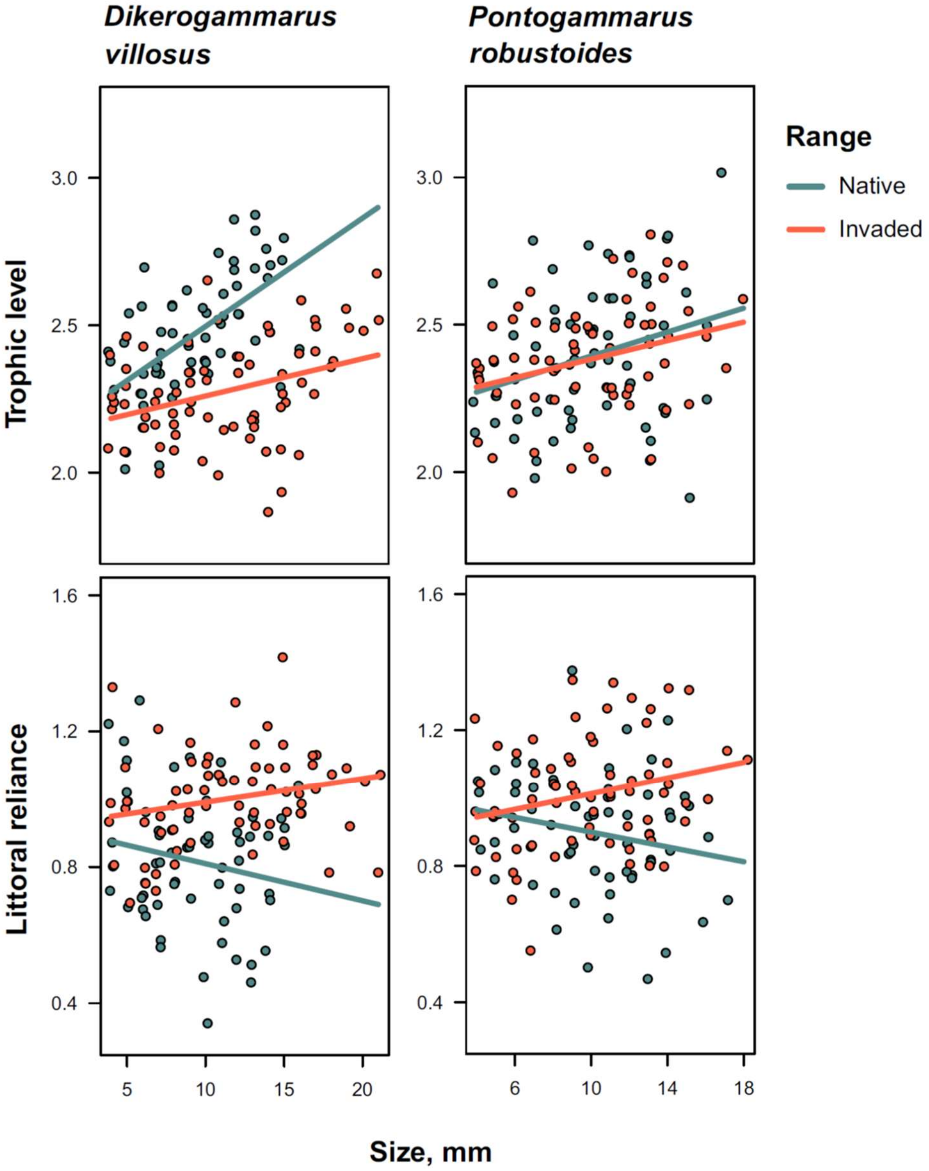


Fig. S3. Size and Range effects on stable isotope metrics of trophic position – trophic level and littoral reliance – in two Ponto-Caspian amphipods – visualisation of linear mixed effects models within datasets containing both adults and juveniles (see Table 1 for effect tests).

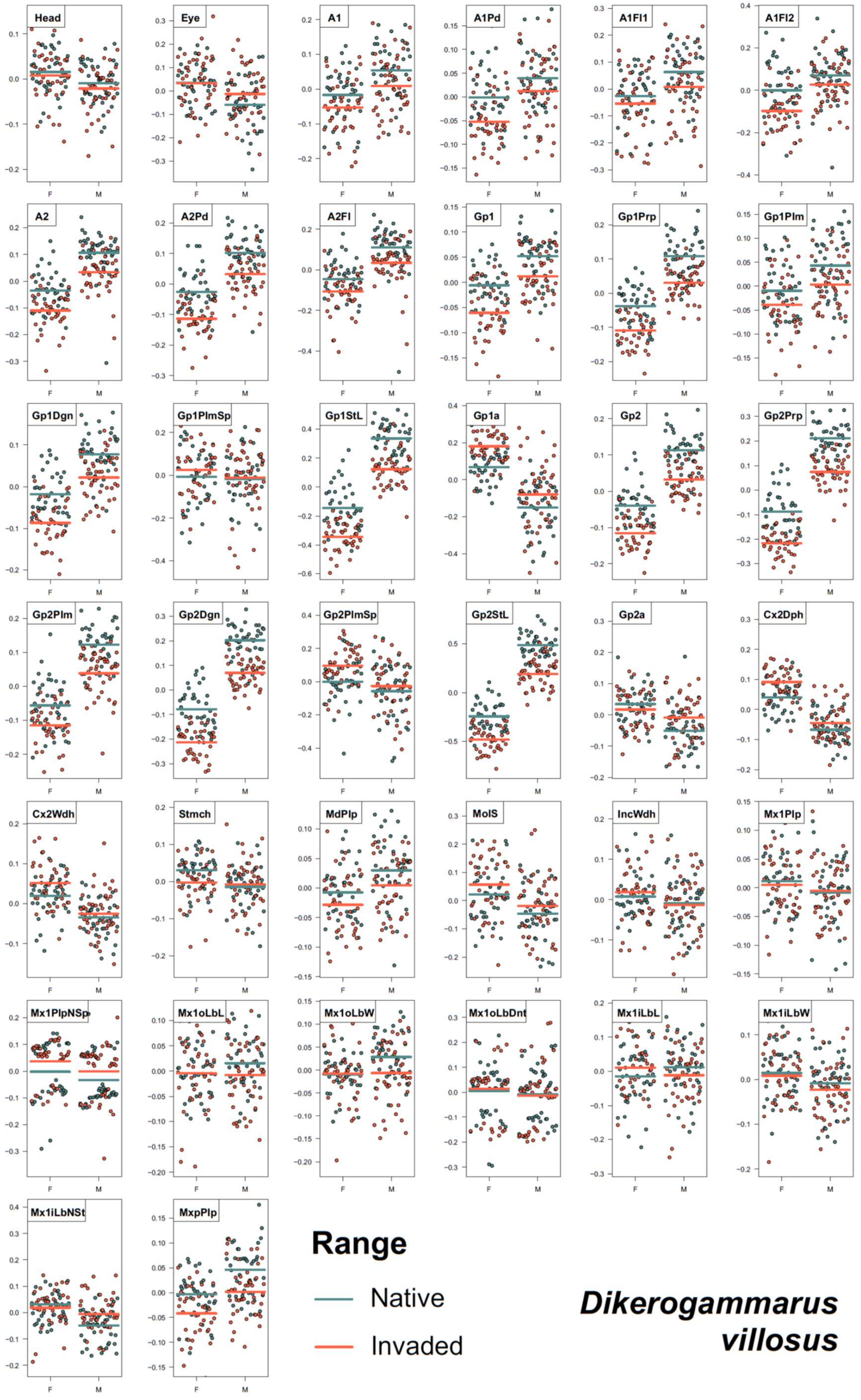


Fig. S4. Sex and Range effects on individual size-corrected morphometric traits in amphipod *Dikerogammarus villosus* – visualisation of linear mixed effects models (see Table S2 for trait definitions and Table S5 for effect tests).

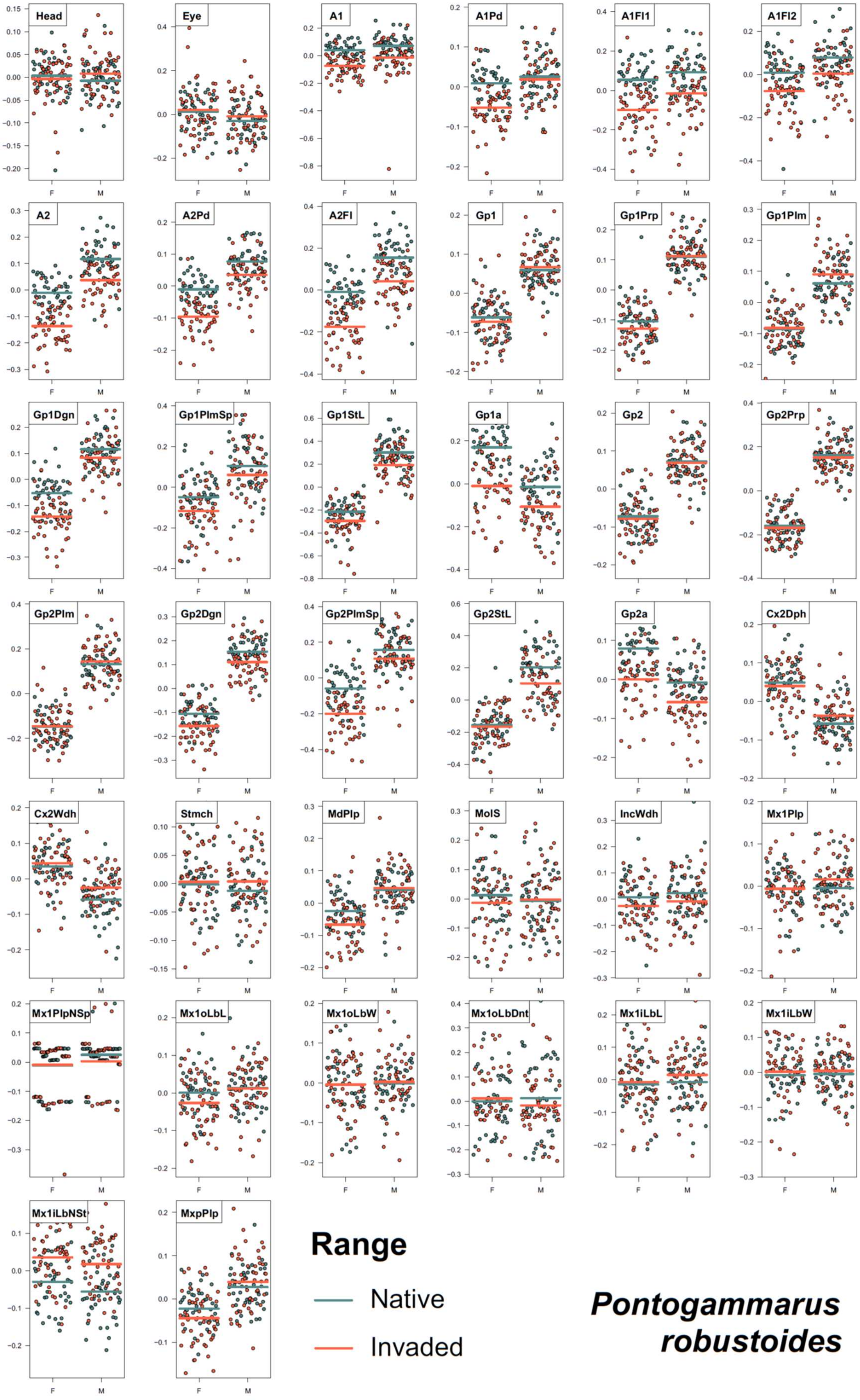


Fig. S5. Sex and Range effects on individual size-corrected morphometric traits in amphipod *Pontogammarus robustoides* – visualisation of linear mixed effects models (see Table S2 for trait definitions and Table S5 for effect tests).

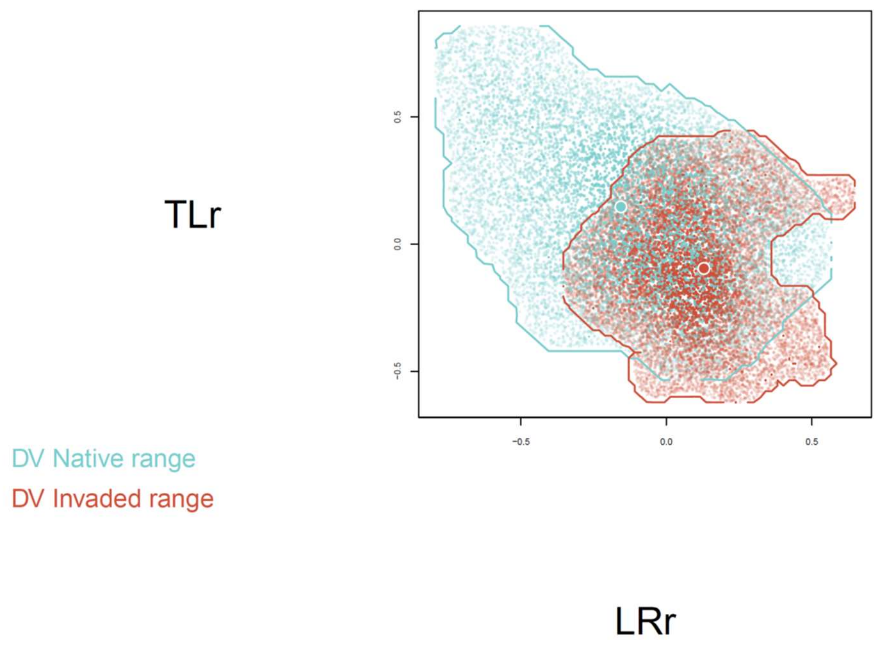


Fig. S6. Stable isotope-derived trophic position hypervolumes for the native and invaded ranges of combined males, females, and juveniles of *D. villosus*. The large dots represent niche centroids. The small dots are 10 000 random points sampled from each hypervolume to delineate its shape and boundary (TLr – residual trophic level, LRr – residual littoral reliance).

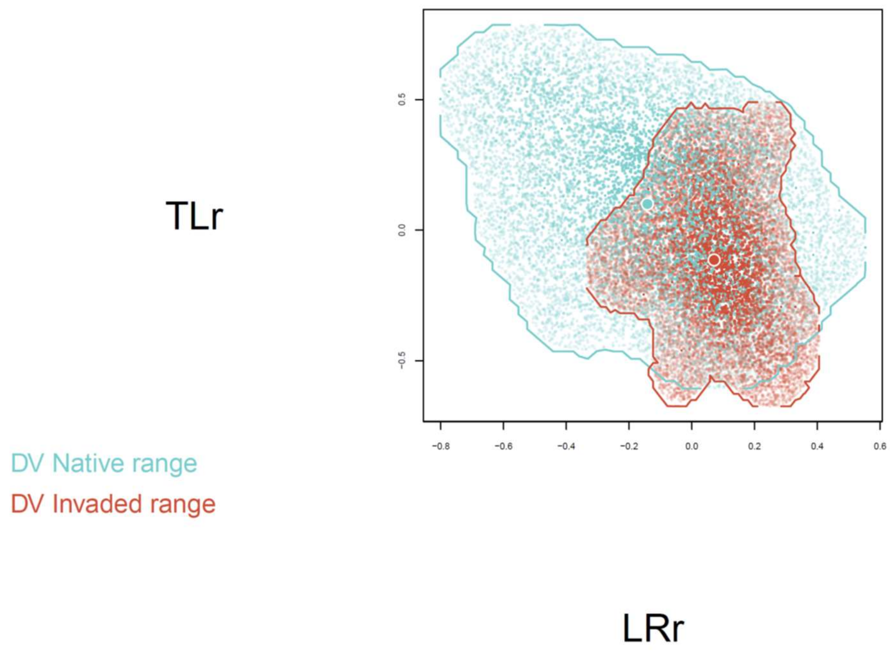


Fig. S7. Stable isotope-derived trophic position hypervolumes for the native and invaded ranges of *D. villosus* males. The large dots represent niche centroids. The small dots are 10.000 random points sampled from each hypervolume to delineate its shape and boundary (TLr – residual trophic level, LRr – residual littoral reliance).

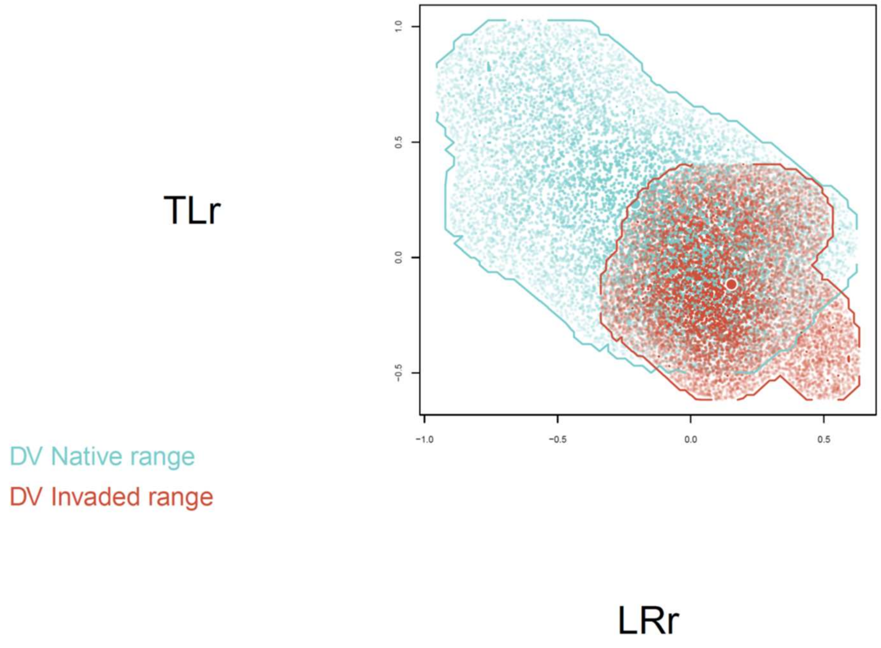


Fig. S8. Stable isotope-derived trophic position hypervolumes for the native and invaded ranges of *D. villosus* females. The large dots represent niche centroids. The small dots are 10.000 random points sampled from each hypervolume to delineate its shape and boundary (TLr – residual trophic level, LRr – residual littoral reliance).

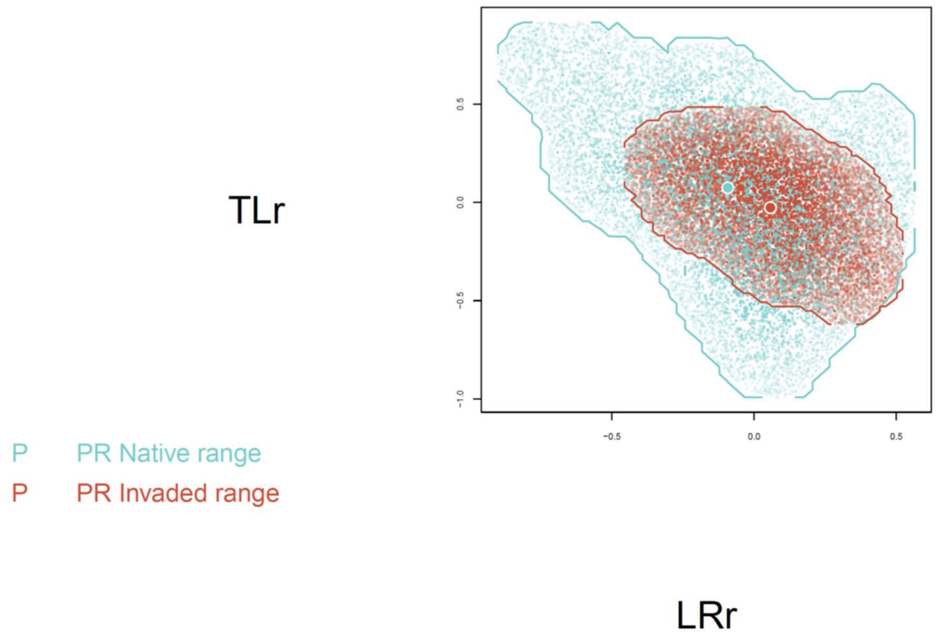


Fig. S9. Stable isotope-derived trophic position hypervolumes for the native and invaded ranges of combined males, females, and juveniles of *P. robustoides*. The large dots represent niche centroids. The small dots are 10.000 random points sampled from each hypervolume to delineate its shape and boundary (TLr – residual trophic level, LRr – residual littoral reliance).

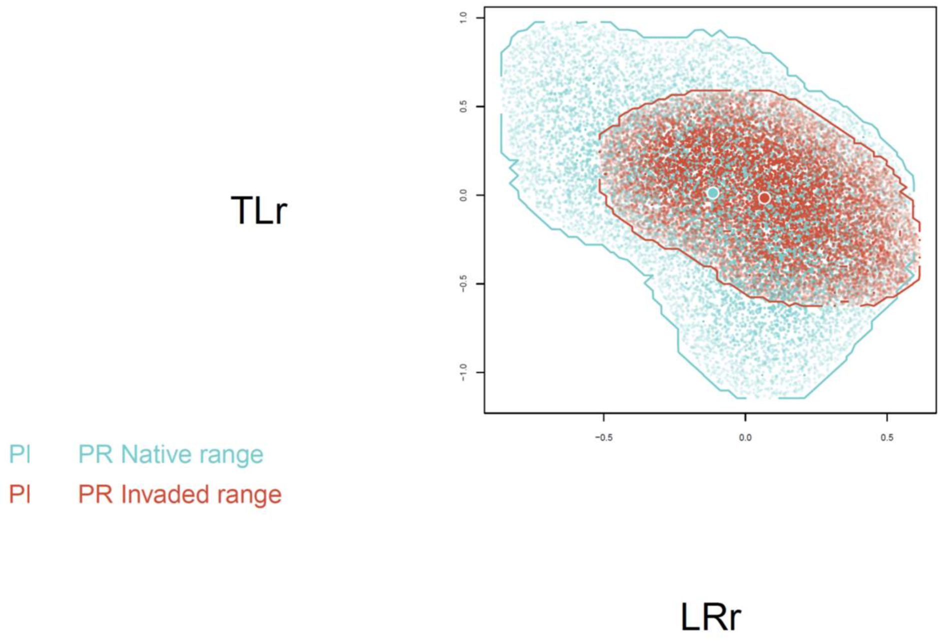


Fig. S10. Stable isotope-derived trophic position hypervolumes for the native and invaded ranges of *P. robustoides* males. The large dots represent niche centroids. The small dots are 10.000 random points sampled from each hypervolume to delineate its shape and boundary (TLr – residual trophic level, LRr – residual littoral reliance).

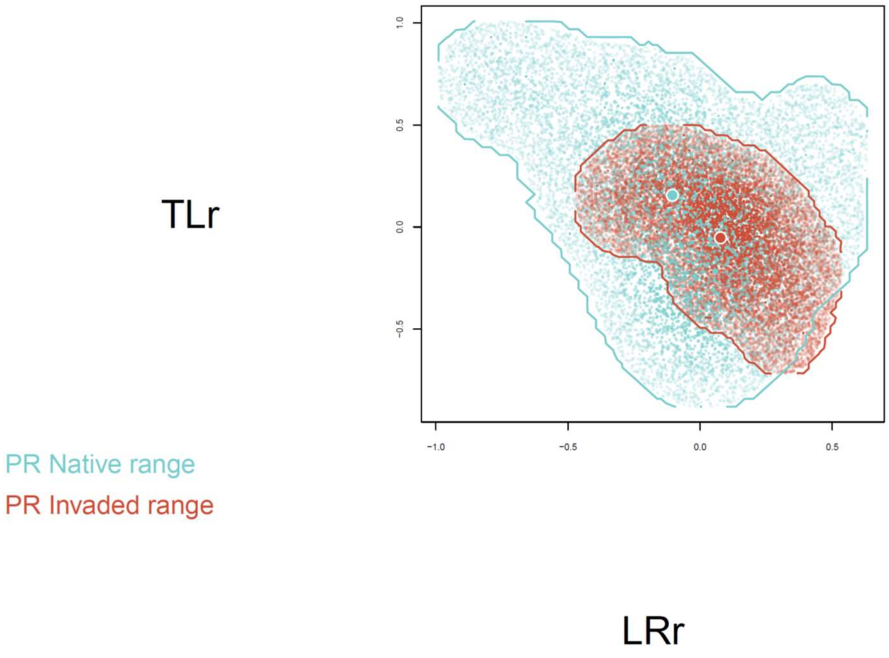


Fig. S11. Stable isotope-derived trophic position hypervolumes for the native and invaded ranges of *P. robustoides* females. The large dots represent niche centroids. The small dots are 10.000 random points sampled from each hypervolume to delineate its shape and boundary (TLr – residual trophic level, LRr – residual littoral reliance).

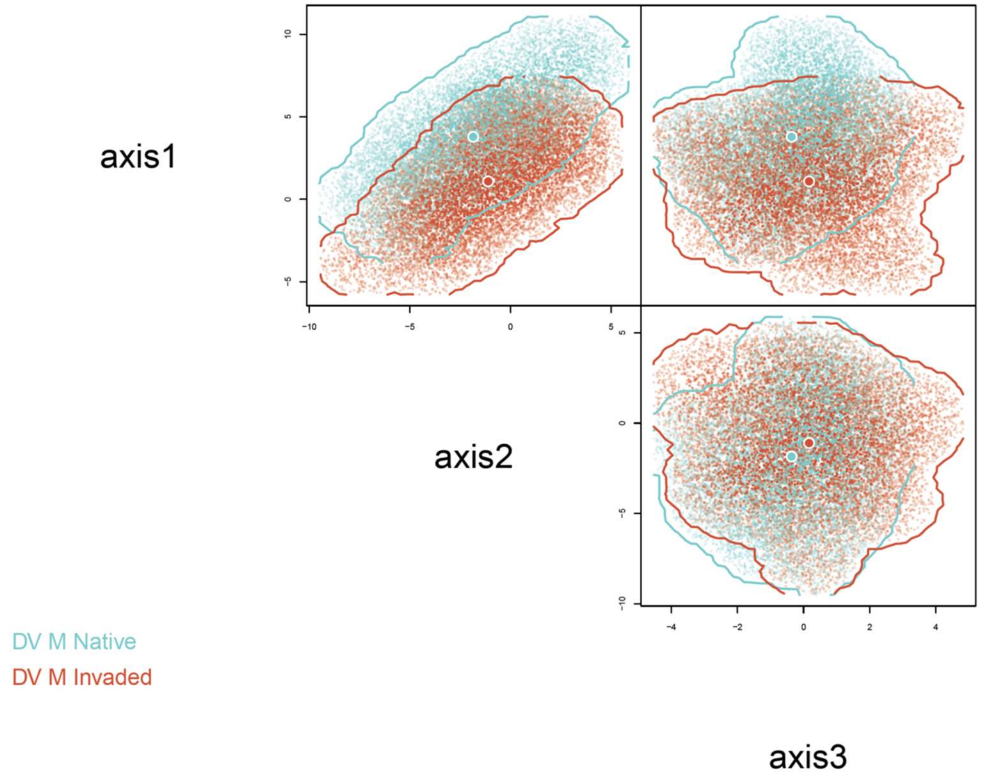


Fig. S12. Morphology hypervolumes for the native and invaded ranges of *D. villosus* males. The large dots represent niche centroids. The small dots are 10.000 random points sampled from each hypervolume to delineate its shape and boundary.

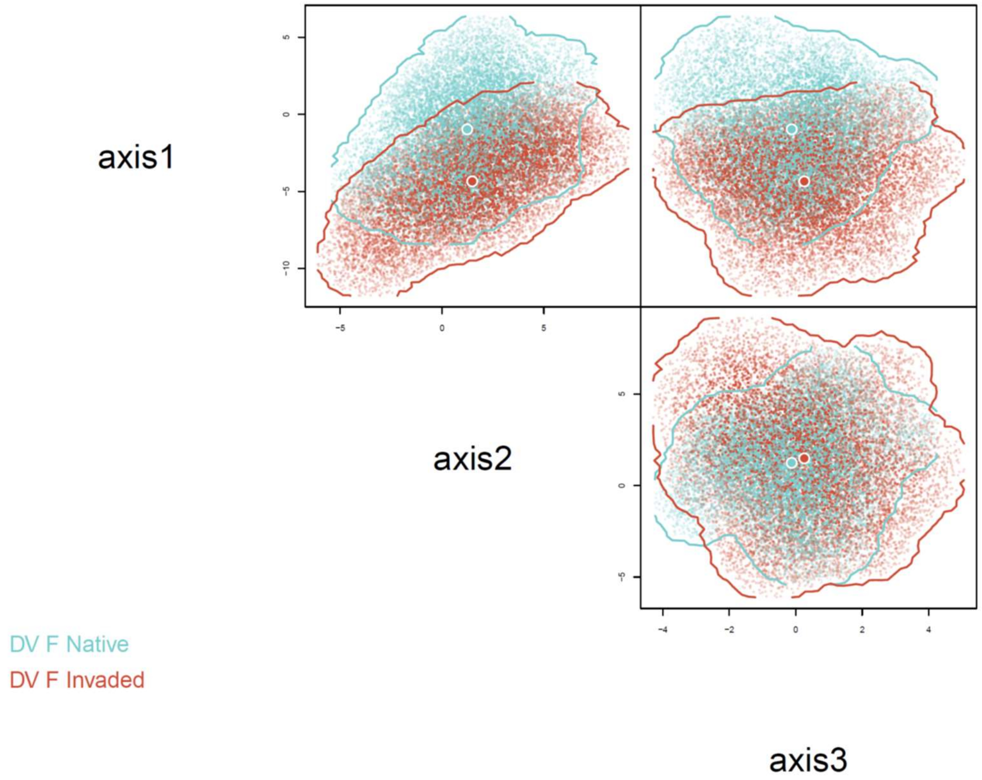


Fig. S13. Morphology hypervolumes for the native and invaded ranges of *D. villosus* females. The large dots represent niche centroids. The small dots are 10.000 random points sampled from each hypervolume to delineate its shape and boundary.

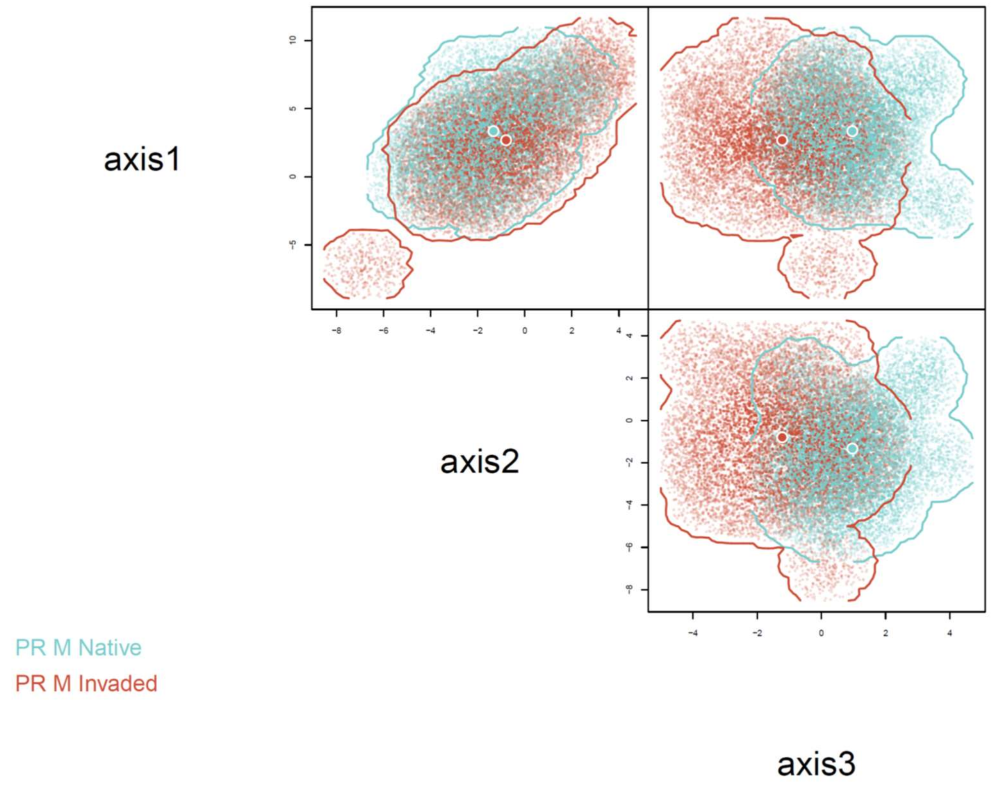


Fig. S14. Morphology hypervolumes for the native and invaded ranges of *P. robustoides* males. The large dots represent niche centroids. The small dots are 10.000 random points sampled from each hypervolume to delineate its shape and boundary.

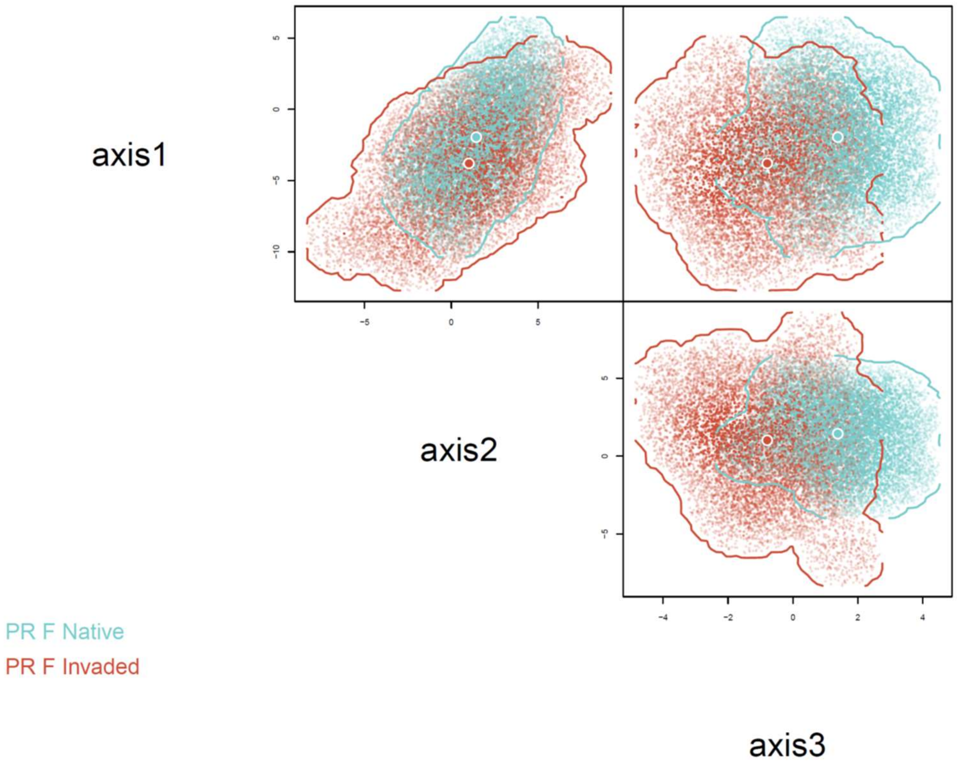


Fig. S15. Morphology hypervolumes for the native and invaded ranges of *P. robustoides* females. The large dots represent niche centroids. The small dots are 10.000 random points sampled from each hypervolume to delineate its shape and boundary.
